# Reading eye movements in traumatic brain injury

**DOI:** 10.1101/496752

**Authors:** Ashwini V C Reddy, Revathy Mani, Ambika Selvakumar, Jameel Rizwana Hussaindeen

**Author notes:** **Corresponding author name and e-mail address:** Jameel Rizwana Hussaindeen.

## Abstract

**Background:** The aim of our study was to measure the reading eye movements in subjects with traumatic brain injury using ReadAlyzer. ReadAlyzer is an objective eye movement recording device that tracks the eye movements while reading.

**Methods:** Reading eye movements were measured using ReadAlyzer in 30 subjects with traumatic brain injury (mild, moderate and severe) who had binocular vision and reading related symptoms and 60 asymptomatic controls.

**Results:** There was a significant decrease in reading eye movement parameters in subjects with traumatic brain injury compared to controls. Reading eye movement parameters were represented in median (IQR). Subjects with traumatic brain injury presented with an increased number of fixations/100 words: 137 (106-159) and regressions/100 words: 24 (12-36), and reduced reading rate 154 (128-173) words per minute. They also had a lesser grade level equivalent: 4.0 (3.0-7.0) and reduced comprehension: 70 (60-80) percentage (Mann-Whitney U test, p<0.05). Reading eye movement parameters were significantly affected in mild and moderate-severe traumatic brain injury subjects compared to controls (Kruskal-Wallis test, p<0.05).

**Conclusion:** Reading eye movement performance using ReadAlyzer was found to be decreased in traumatic brain injury. Reading assessment may serve as a clinical measure to understand the oculomotor system due to traumatic brain injury.

---

Reading is one of the most important visual functions in daily living. The act of reading is highly complex involving an integrated function of oculomotor, sensory, cognitive, and attentional aspects.^1^ Oculomotor system primarily involves execution of vergence, versions and accommodation during fixations, reading, writing and while viewing any target in the environment.

A normal reading is comprised of accurate, rhythmical and spontaneously executed sequences of saccadic eye movements interspersed with brief fixational pauses.^1,2^ Reading related saccadic eye movements are 1-3 degrees in amplitude and the saccadic latencies are 30-60 msec.^2^ The presence of accurate saccadic tracking, synchronised ocular accommodation and vergence is required for efficient reading.

In traumatic brain injury (TBI), multiple brain areas and their functions are adversely affected because of the diffuse axonal injury (DAI). A physical damage to the underlying neurons, such as stretching, twisting, and shearing of the neurons can cause an impairment resulting in a range of sensory, oculomotor, perceptual and structural abnormalities.^1,3^

Symptoms following TBI may persist for seconds to minutes after the event and usually resolve within 12 weeks but may continue for months or even years.^4^ Impairment of the oculomotor subsystem following TBI, also adversely affects the naturalistic pattern of reading. Ninety per cent of the visually symptomatic mild TBI (mTBI) group exhibited oculomotor dysfunction (OMD) following the head trauma.^3^

Reading eye movements are one among the important oculomotor functions that enable an individual to read or comprehend a paragraph using basic oculomotor functions. Studies have shown impaired reading eye movement parameters due to head injuries. Thiagarajan et al., had investigated reading eye movements in mild TBI using Visagraph and found that the subjects had significantly reduced reading rate, an increased number of fixations/100words, a higher number of regressions/100words, and decreased grade-level efficiency.^1^ During reading, an individual with TBI exhibits hypometric saccades (<1-degree amplitude) and increased saccadic latencies (>200 msec).^2^ Considering the extensive neural network of the oculomotor subsystems, a global damage in TBI could compromise precise oculomotor control, leading to reading dysfunction and an unreceptive quality of life (QoL).^1^

The assessment of reading eye movements is highlighted in this study because eye movements are considered as novel visual biomarkers to predict the high-risk population from persisting with symptoms of TBI.^5,6^ There is a limited literature on clinically-based evaluation of reading eye movement parameters with objective eye movement recordings for these individuals in India. Understanding the pattern of reading eye movements is essential as eye movements are deliberated to be one of the key elements in assessing the functional integrity of the brain. This assessment can potentially support early visual intervention in reading dysfunction. Therefore, we present our study that investigated the impact of TBI on reading eye movements using ReadAlyzer, an objective eye movement recording device.

## METHODS

### Study Design

A prospective comparative study was conducted between April 2015 and February 2016 in the Neuro-Optometry Clinic at a tertiary eye care center, India. The study adhered to the tenets of Declaration of Helsinki and the investigational procedures were reviewed and accepted by the Institutional Review Board and Medical Ethics Committee.

### Subjects

Thirty subjects with TBI and 60 controls were included in the study. The sample size was estimated as 30 subjects diagnosed with TBI and 60 age-matched controls considering a 1:2 ratio between the cases and the controls. Subjects with TBI were referred from the Neuro-ophthalmology department if they complained about any one of the symptoms of reading difficulty, headache, eye strain, dizziness. Age-matched subjects who volunteered to participate in the study were chosen as controls. Inclusion and exclusion criteria for the cases and the controls are presented in Table 1. A duly signed, written informed consent was obtained from all the study participants. All the subjects received a comprehensive eye examination which included history taking, refraction, pupillary evaluation, extraocular motility, anterior and posterior segment examination. This was followed by a detailed neuro-optometric evaluation.

**Table 1:** Inclusion and Exclusion Criteria for study subjects.

| Subjects | Inclusion Criteria | Exclusion Criteria |
| --- | --- | --- |
| **Cases** | • Age range: 18 – 60 years<br>• Best corrected visual acuity ≥ 6/9 for distance and N6 for near in worse eye<br>• TBI cases with one or more visual symptoms (For example: a headache, skipping of lines while reading, blur, eye strain) and one clinical sign (For example: receded near point of convergence) of oculomotor or non-strabismic binocular vision anomalies<sup>3</sup><br>• Onset and persistence of visual symptoms at least six months' post-injury<sup>1</sup><br>• Ability to understand the test instructions<br>• Intact visual field with Confrontation and Amsler test<br>• Proficiency with the English language<sup>†</sup><br>• Stable systemic conditions for 5 years (For example: Diabetes Mellitus & Hypertension under control) | • Central or paracentral visual field defects with Confrontation test/Humphrey visual field test that hinder reading performance<br>• Constant strabismus, amblyopia, nystagmus, an ocular disease in either eye (For example: Glaucoma) |
| **Controls** | • Age range: 18 – 60 years<br>• Proficiency with the English language<sup>†</sup><br>• Normal binocular vision parameters<br>• Non-symptomatic for reading or near work<br>• Stable systemic conditions (For example: Diabetes Mellitus & Hypertension under control) | • Best corrected visual acuity < 6/9 for distance and < N6 for near in either eye<br>• Constant strabismus, amblyopia, nystagmus, an ocular disease in either eye (For example: Glaucoma) |

<sup>†</sup> Proficiency with the English language was set as an inclusion criterion as study participants were asked to read English passages using ReadAlyzer

### Testing procedures

#### Neuro-optometric examination

A detailed history of the nature of injury and symptoms during the post-injury period were obtained from the TBI subjects. At the time of recruitment, subjects with TBI were classified into mild, moderate, and severe grades based on the Glasgow Coma Scale (GCS), post-traumatic amnesia (PTA) and loss of consciousness (LOC) reported either in the records of emergency department or hospital discharge summary or by iterative questioning about the traumatic event to the subject or subject’s caretaker. GCS is a 3- to 15-point scale used to assess a patient’s level of consciousness and neurologic functioning; scoring is based on motor, verbal, and ocular responses. A score between 13-15 is mild, 9-12 is moderate and 3-8 is severe. PTA is the time elapsed from injury to the moment when patients can demonstrate continuous memory of what is happening around them. PTA < 1day is mild, 1-7 days is moderate and >7 days is severe. Duration of loss of consciousness is classified as mild (LOC < 30 min), moderate (LOC 30 min to 6 hr.), or severe (LOC >6 hr.). ^7,8^ In most cases, GCS, PTA and LOC were obtained at the time of admission to the hospital or from the records of discharge summary and in some cases, the GCS scale was used to probe the events that occurred during the injury.

#### Reading eye movement assessment

Reading eye movements were assessed objectively using ReadAlyzer™ (Compevo AB, Markvardsgatan, Stockholm, Sweden). ReadAlyzer consists of infra-red emitters and detectors mounted in a safety goggle. It can determine the eye positions by sensing several infrared reflections from the cornea. The measuring speed of the instrument is 60 Hz with a better angular resolution compared to Visagraph II. Head movements are automatically compensated for analysis by the ReadAlyzer software. ^9-11^ Subject wore the eye movement goggles and the near interpupillary distance was adjusted. The test paragraphs were placed 40 centimetres from the corneal plane or habitual correction centred along the subject’s midline.

#### Reading test

Eye movements were recorded while the subject read a short English paragraph silently. The highest-grade level paragraph (Grade 10 – for adults) was used for measurement. There were five different passages in Grade 10. The subject read one practice paragraph following which two trials were made with different passages. The second trial was taken as the final reading to assure a stable baseline measurement.^12^ A comprehension test comprising 10 “yes” or “no” responses were also administered to confirm the subject’s comprehension. After the recording, the system performed an automatic analysis and provided a report in a ***Reading Profile*** format (Figure 1). Reading parameters included fixations per 100 words (progressive saccades), regressions per 100 words (backward saccades), fixation duration (sec) which is the average length of time (in parts of a second) the eyes paused or fixated, reading rate (words per minute), grade level equivalent (GLE) which is the weighted average of the grade levels for the subject’s fixations, regressions and reading rate yielding a combined grade level, and comprehension (%) which is percentage of correct answers. Seventy per cent or more was acceptable. There are also large right-to-left oblique saccadic eye movements called saccades in return-sweep which occur when one must shift to the next line of print.^11^ Age-matched controls with normal binocular vision parameters were administered with ReadAlyzer test.

**Figure 1:**
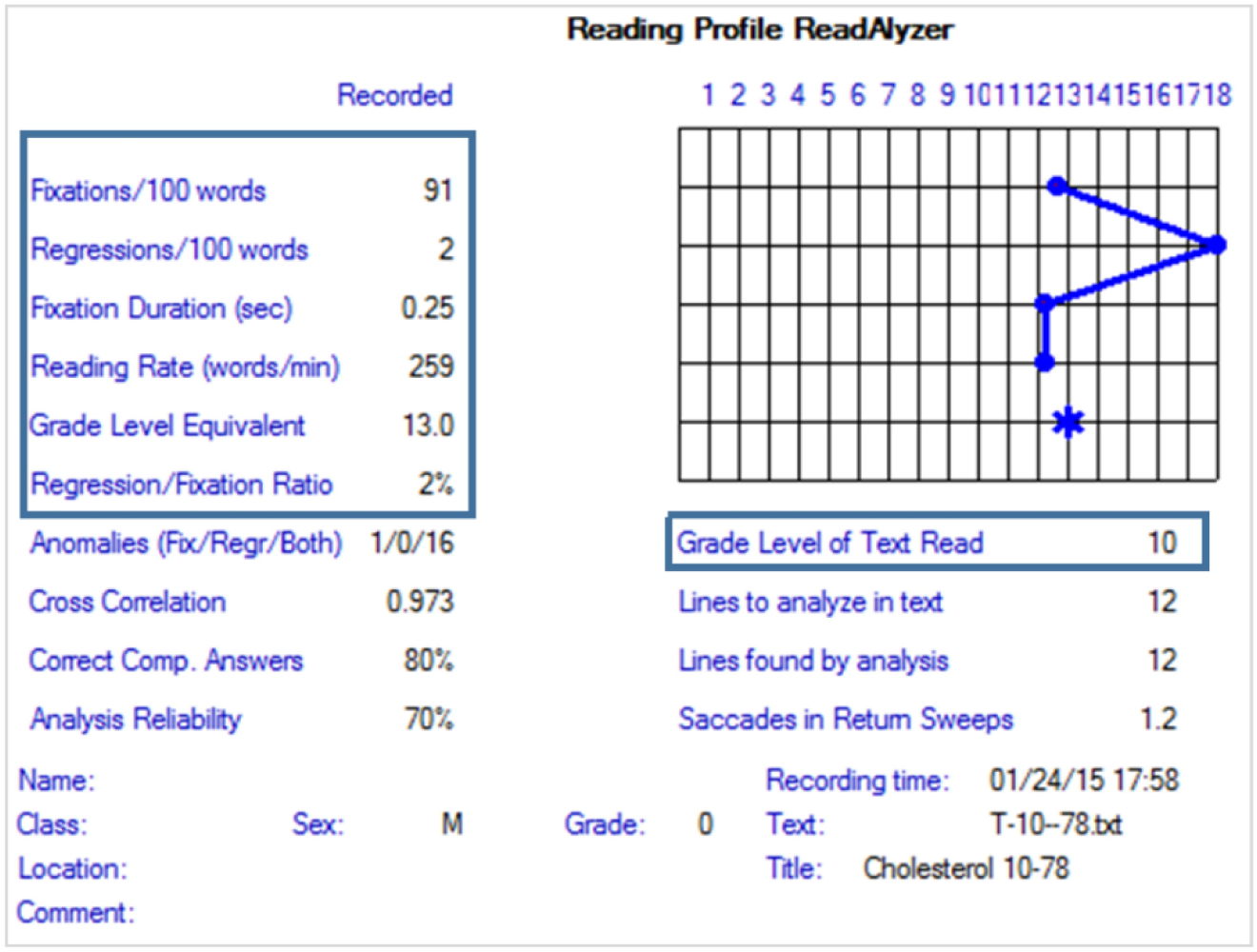
Reading Profile of a normal subject recorded with ReadAlyzer (Report taken from ReadAlyzer™)

### Statistical Analysis

Clinical details of the study participants were entered in Microsoft Excel 2013 and statistical analyses were performed using SPSS (Statistical Package for Social Sciences, Version 17.0, SPSS Inc., Chicago). Non-parametric tests were done as the data did not follow normality (Shapiro-Wilk test). Appropriate coding was generated for categorical variables. Mann-Whitney U test was used to compare the values between TBI cases and controls. Kruskal-Wallis test was used to compare the values between different grades of TBI with controls. As the moderate and severe TBI groups had a lesser sample size, these two groups were combined as MS-TBI for analysis. Spearman’s correlation was used to understand the relationship between variables. Median and interquartile range values were used to represent the data. The alpha error was set as 5%.

## RESULTS

Ninety subjects (30 cases and 60 controls) were included for statistical analysis. The mean age ± SD of the TBI and controls was 28.7 ± 8.5 years (18.4 - 58.9) and 28.4 ± 7.7 years (20.4 - 57.0) respectively. The difference in age was not statistically significant between the two groups *(Chi-square test, p=0.052)*. There were 18 mild TBI (mTBI) and 12 moderate-severe (MS-TBI) (4 moderate and 8 severe) cases of TBI.

### Aetiologies of TBI

In the present study, road traffic accidents (RTA) (n=24, 80%) was the most common cause of TBI followed by hit (n=4, 13%) and fall from height (n=2, 7%). All RTA’s were caused merely due to two-wheelers. Four subjects who had a history of an object striking their head which was defined as hit and two subjects had TBI due to falling from a height. The median (IQR) post-injury periods of mild, moderate and severe TBI were 2 (0.6-5), 1.2 (0.5-5.9), 2.5 (0.7-3.7) years respectively.

### Symptoms of TBI subjects

TBI subjects in the current study self-reported their symptoms which persisted past 6 months from the onset of TBI (Figure 2). In the total TBI sample, reading difficulty (87%) was the most frequent visual issue followed by eye strain (47%), headache (40%), vertigo/dizziness (10%) and double vision (10%). A majority of mTBI subjects reported symptoms of reading difficulty, eyestrain and dizziness, and, MS-TBI subjects had issues such as a headache primarily followed by reading difficulty and eye strain (Table 2).

**Table 2:** Symptoms in mild TBI and moderate-severe TBI.

| Symptoms | mTBI <sup>†</sup><br>n (%) | MS-TBI <sup>‡</sup><br>n (%) |
| --- | --- | --- |
| Reading difficulty | 18 (100) | 9 (75) |
| Eye strain | 14 (77.5) | 10 (83) |
| Headache | 12 (66.7) | 12 (100) |
| Vertigo/Dizziness | 0 | 3 (25) |
| Double vision | 0 | 7 (58) |

<sup>†</sup> mTBI – Mild TBI; <sup>‡</sup> MS-TBI – Moderate-Severe TBI

**Figure 2:**
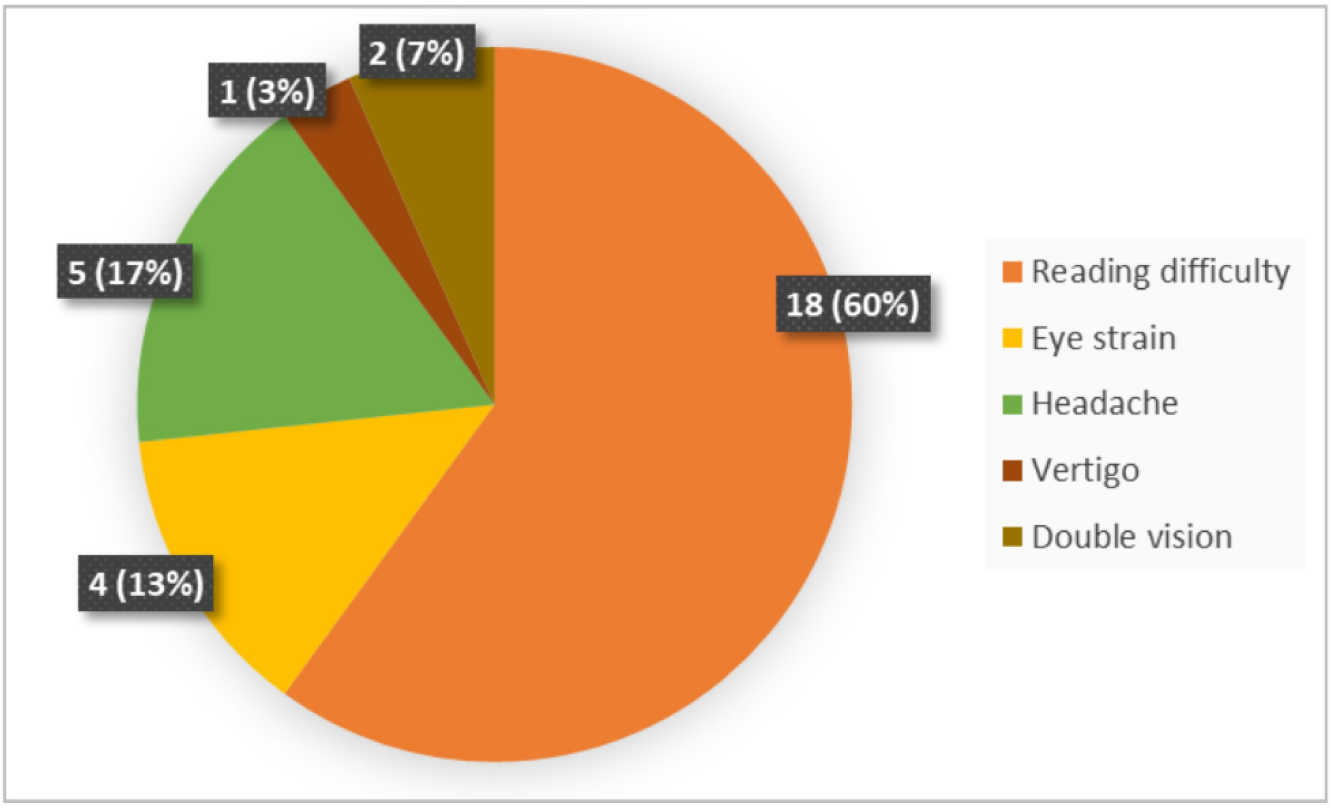
Symptoms of all TBI subjects [n (%)]

### Reading Eye Movement parameters: TBI *vs* Controls

The results of the oculomotor-based reading eye movement assessment using ReadAlyzer™ were compared with age-matched controls (Table 3). Subjects with TBI presented with increased number of fixations/100 words: 137 (106-159), regressions/100 words: 24 (12-36), reduced reading rate of 154 (128-173) words per minute, lesser comprehension: 70 (60-80) percentage, lower grade level equivalent: 4.0 (3.0-7.0) and increased return sweep saccades: 1.7 (1.2-2.4) *[represented in median (IQR); p <0.01].*

**Table 3:** Comparison of Reading test parameters obtained from ReadAlyzer between controls and TBI.

| Reading test parameters | TBI † | Controls | $p^*$ |
| --- | --- | --- | --- |
|  | Median (IQR) [n=30] | Median (IQR) [n=60] |  |
| Fixations/100 words (No.) | 137 (106-159) | 92 (76-102) | <0.001 |
| Regressions/100 words (No.) | 24 (12-36) | 10 (7-14) | <0.001 |
| Fixation Duration (sec) | 0.30 (0.27-0.33) | 0.29 (0.27-0.32) | 0.2 |
| Reading rate (words per min) | 154 (128-173) | 214 (199-244) | <0.001 |
| Comprehension (%) | 70 (60-80) | 90 (70-100) | 0.008 |
| Grade Level Equivalent (GLE) | 4.0 (3.0-7.0) | 10.0 (8.0-12.0) | <0.001 |
| Saccades in Return Sweeps (No.) | 1.7 (1.2-2.4) | 1.5 (1.3-1.8) | 0.022 |

\* Mann-Whitney U test; $p$ value represents the statistical significance for the comparison between TBI and control groups
† TBI – Traumatic Brain Injury

To understand the reading eye movements based on the severity of TBI, a comparison between three groups (controls, mTBI and MS-TBI) was conducted which showed a significant difference between the three groups *(p<0.01)* (Table 4). Post hoc analysis revealed a significant difference between controls and mTBI (p<0.01), controls and MS-TBI (p<0.01) and no statistically significant difference was noted between mTBI and MS-TBI (p=0.43).

**Table 4:**
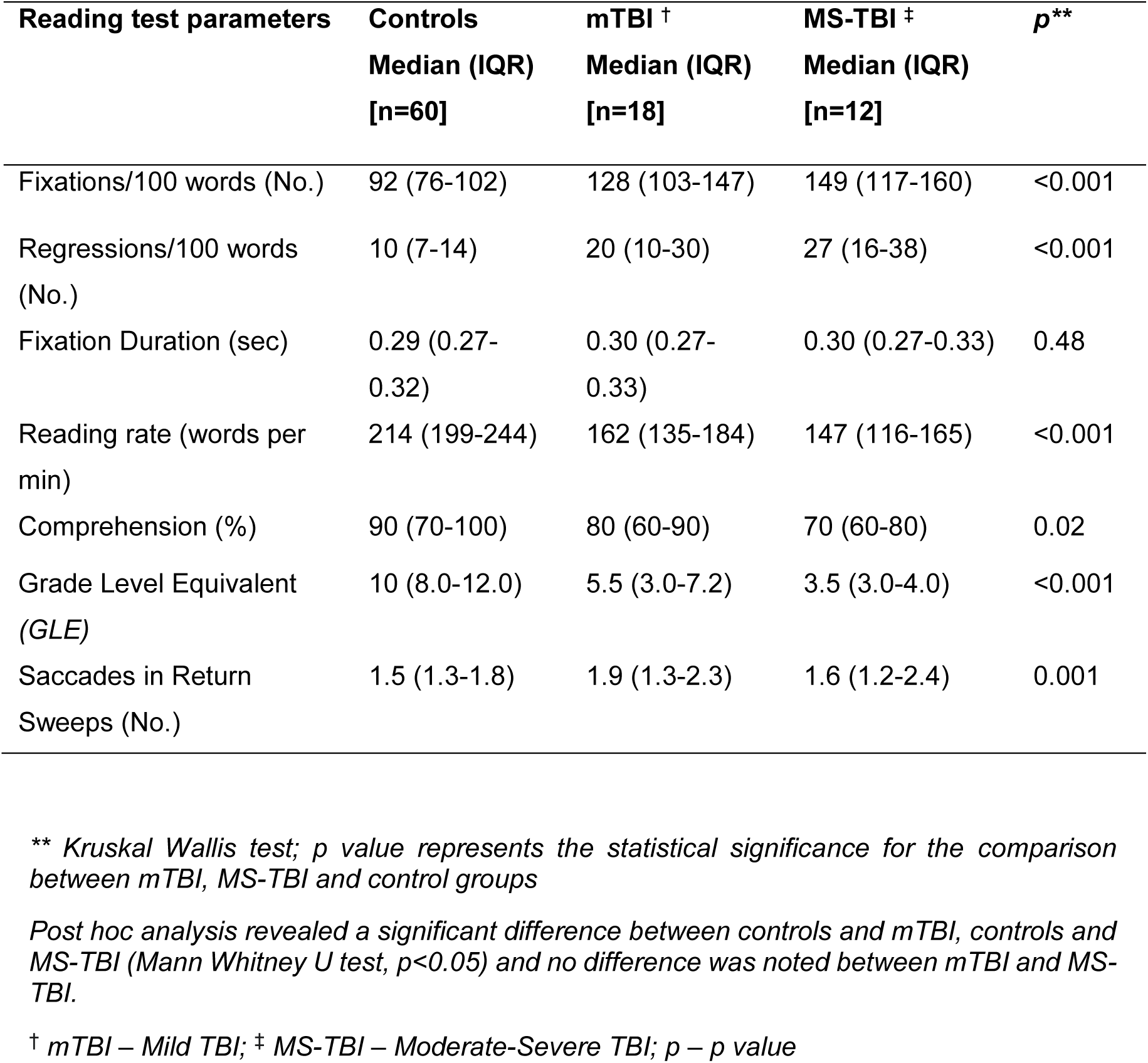
Comparison of Reading test parameters obtained from ReadAlyzer between controls, mTBI and MS-TBI.

### Correlation between number of fixations per line and reading rate in TBI

The relationship between the number of fixations/100 words and reading rate in TBI subjects showed a significantly strong negative correlation (Figure 3) (Spearman’s correlation, *r = −0.823, n=30, p= <0.001*) in subjects with TBI.

**Figure 3:**
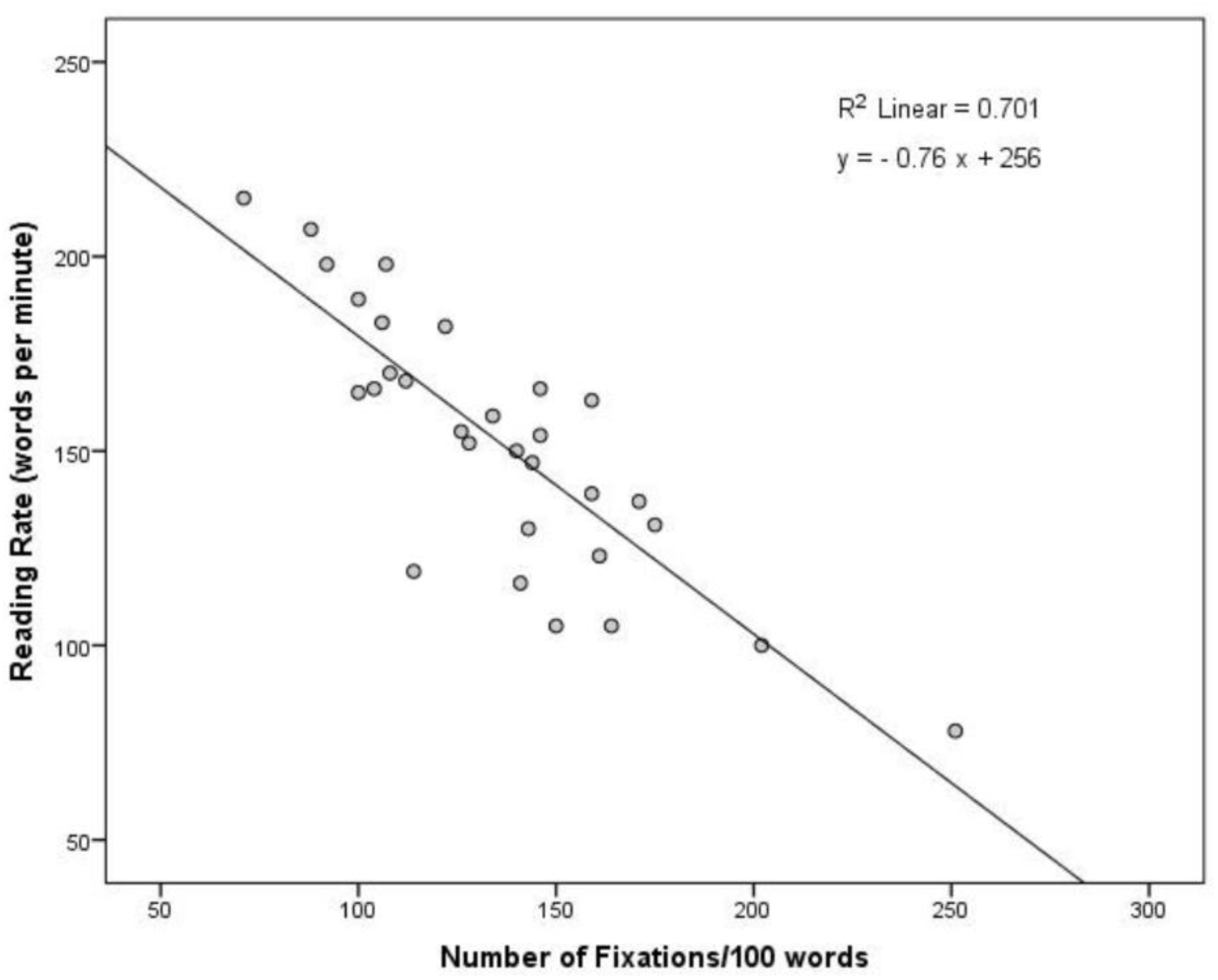
Correlation between number of fixations per line and reading rate in TBI

## DISCUSSION

In our study, reading eye movement parameters in subjects with TBI were evaluated and compared with age-matched controls. The current research is the first to study and report eye movement parameters during reading in TBI in India.

It is important to address the physical and visual issues following TBI as it can result in morbidity, mortality, disability and socioeconomic losses in many developing countries.^13^ In India, an assessment of injury pattern of RTAs that had collisions tangled with head injuries was caused frequently by two-wheelers (62%) and less likely by four-wheelers (12%);^14^ whereas, in western countries, most of the accidents were found due to four-wheelers (79%). In the present study, twenty-four subjects (80%) reported RTA due to two-wheelers and were diagnosed to have TBI. This scenario is due to the unprecedented motorization and unacceptance of safety policies among the two-wheeler drivers.^14,15^

Reading is an essential task in every individual’s life. Any mishap due to TBI can affect the oculomotor system resulting in reading difficulty and affect the quality of life (QoL). ^16,17^ There were scenarios in the current study were subjects with TBI (60%) gave up their regular reading habits due to troubling eye-related symptoms. Therefore, we studied reading eye movement parameters in subjects with TBI and the results from our study showed that eye movements are affected. This, in turn, affected the naturalistic reading ability of a compromised individual compared to a normal.

The evaluation of clinically-based reading eye movements has provided insight into the functional integrity of the brain. ReadAlyzer™ was used to evaluate the reading eye movements. It has been a valid, clinical tool that provided consistent, objective and automated results on reading eye movement parameters. The advantage of this instrument is that the infrared cameras have allowed real-time observation of eye movements during recording. Dynamics of saccades such as saccadic latency and accuracy are also known to be affected by ageing.^18^ Therefore, study sample recruitment was done by ensuring that controls were age-matched to a TBI subject. We also correlated reading eye movement parameters with age, but results did not reveal any significant correlation with age.

For a subject with normal visual function, based on their grade level, the expected reading rate is 250-280 words per minute with 90 fixations per 100 words and 15 regressions per 100 words according to Taylor’s normative data for the adult American population.^2^ In the present study, controls also had a lesser reading rate: 214 (199-244) words per minute compared to an established Taylor’s normative data. These differences suggested that reading an English text is based on the familiarity with language and vocabulary. ^2^ As English is a second language in India, the fluency and speed of reading are variable when compared to native English speakers. Hence, reading eye movement parameters of TBI subjects were assessed by comparing with age-matched controls due to the lack of evident age-based normal reading rate for our population. All the subjects (TBI and controls) in the present study were ensured that they held a basic degree with fluency in English. Individuals with TBI in the present study demonstrated significantly reduced reading rate, increased number of fixations, and a higher number of regressions. The results suggested that subjects with TBI had a low degree of saccade automation, and they resulted in making an excessive number of unwanted saccades which reflected in their reading. Studies explained that the low gain in the saccadic amplitude of the primary saccade resulted in a hypometric saccade. Therefore, a corrective subsequent saccade was made to achieve the anticipated saccadic amplitude.^1,19^ These corrective saccades resulted in an increased number of fixations and regressions with poor reading eye movements. Subjects with TBI also had reduced comprehension which revealed a problem with inference and short-term memory in answering the questions along with basic demands of oculomotor coordination compared to controls.^11^ With all these parameters being reduced, the grade level equivalent was also lesser in subjects with TBI, as they read 5 grade levels lesser than controls. This finding of an increased number of inaccurate reading eye movements is consistent with a study reported by Thiagarajan, et al. on mTBI population which were measured using Visagraph (2014). ^1^ It was reported that during reading, an individual with TBI exhibits hypometric saccades (<1 degree in amplitude), ^1,20^ increased saccadic latencies (>200 msec), increased number of fixations (>90 per 100 words), regressions (>12 per 100 words) and reduced reading rate (<250 words per minute).^1^

The information on clinically-based reading eye movements when translated into the natural reading process helped us to interpret reading dysfunction. Comparison of reading eye movement parameters between mild TBI (mTBI) vs. moderate and severe TBI (MS-TBI) with controls highlights that the oculomotor system is compromised both in mTBI and MS-TBI. Mild TBI and MS-TBI did not show any statistically significant difference even though the outcome measures were relatively affected in MS-TBI. Alternatively, the extent of reading dysfunction in TBI might not be truly dependent on injury severity. There lies a possibility that visual functions are vulnerable to damage regardless of the severity of the injury. Similarly, the symptoms of reading difficulty were more profound in mTBI compared to MS-TBI inferencing that mTBI is also affected like MS-TBI. It has been described that the susceptibility of extensive neural networks affected the multiple brain regions associated with control, execution, initiation and generation of saccades leading to reading dysfunction. ^1, 3^

## CONCLUSION

This study adds evidence to the impaired reading eye movement performance in TBI invariable to the severity. ReadAlyzer, being a simple instrument has helped us to understand the quantitative reading parameters in TBI. It has been highlighted in the present study that reading is affected in all severity of TBI. The degree of reading impairment increased with the severity of the injury as an important clinical finding. These clinically-based reading eye movements were addressed previously in mTBI, but not in MS-TBI using ReadAlyzer. Oculomotor testing is thus sensitive to detect subtle defects in all grades of TBI.

Having understood about the visual sequelae in TBI, it is also important to rehabilitate these subjects with oculomotor vision therapy. Studies have shown that oculomotor rehabilitation can significantly improve overall reading and result in behavioural changes with a progressive effect on the QoL. ^1,20-22^ This improvement has also been observed in a case of mTBI with convergence insufficiency and reading dysfunction that we reported. ^23^ Neuro-optometric vision therapy facilitated the subject to recuperate from the compromised state and perform better in his daily living activities.

The limitations of the study include inadequate sample size in moderate and severe TBI groups. Visual symptoms were not quantified using a validated questionnaire used for TBI. Subjects with English language proficiency were only used as the reading passages were in English. Test paragraph with different regional languages that match the corresponding grade level equivalent may serve for non-English proficiency subjects.

An extensive future research in the objective assessment of eye movements in TBI may help neuro-optometrists to understand the occurrence of deficits in reading eye movements. This may also help the clinicians to evaluate the reading deficits in the regular neuro-optometric work up and also to monitor recovery/improvement with intensive neuro-optometric intervention.

## ACKNOWLEDGEMENTS

The authors would like to acknowledge Mr Kurt Mirdell, Developer of Visagraph II and ReadAlyzer for answering technical queries.

